# The Latent Genetic Structure of Impulsivity and its Relation to Internalizing Psychopathology

**DOI:** 10.1101/2020.01.07.897231

**Authors:** Daniel E. Gustavson, Naomi P. Friedman, Pierre Fontanillas, Sarah L. Elson, the 23andMe Research Team, Abraham A. Palmer, Sandra Sanchez-Roige

## Abstract

Factor analyses suggest that impulsivity traits that capture tendencies to act prematurely or take risks tap partially distinct constructs. We applied genomic structure equation modeling to evaluate the genetic factor structure of two well-established impulsivity questionnaires, using published genome-wide association study statistics from up to 22,861 participants. We also tested the hypotheses that delay discounting would be genetically separable from other impulsivity factors, and that emotionally-triggered facets of impulsivity (urgency) would be those most strongly genetically correlated with an internalizing latent factor. A five-factor model best fit the impulsivity data. Delay discounting was genetically distinct from these five factors. As expected, the two urgency subscales were most strongly related to an Internalizing Psychopathology latent factor. These findings provide empirical genetic evidence that impulsivity can be disarticulated into distinct categories of differential relevance for internalizing psychopathology. They also demonstrate how measured genetic markers can be used to inform theories of psychology/personality.

## Introduction

Impulsivity is a construct common to many theories of personality (Evenden, 1999; Eysenck & Eysenck, 1985; Tellegen, 1982). Impulsive personality traits (**IPTs**) typically refer to a tendency to act without planning or self-control (lack of premeditation), the inability to resist temptations while experiencing positive or negative affect (positive urgency and negative urgency), the inability to persist on difficult tasks (lack of perseverance), or the tendency to enjoy exciting situations (sensation seeking). These five IPTs are measured by the **UPPS-P** Impulsive Behavior Scale (Lynam, Smith, Whiteside, & Cyders, 2006; Whiteside & Lynam, 2001), arguably the most common questionnaire assessing impulsivity. Another widely used instrument is the Barratt Impulsiveness Scale (**BIS**), which also focuses on the tendency to act without premeditation (Barratt, 1993; Patton, Stanford, & Barratt, 1995).

IPTs are phenotypically and neurologically dissociable from one another (Dalley, Everitt, & Robbins, 2011; MacKillop et al., 2016; Reynolds, Ortengren, Richards, & de Wit, 2006) and show divergent associations with psychiatric disorders, particularly substance use and internalizing psychopathology (Cyders & Smith, 2007; Johnson, Carver, & Joormann, 2013). For example, a meta-analysis of UPPS-P (115 studies, 40,432 individuals; Berg, Latzman, Bliwise, & Lilienfeld, 2015) revealed that only three of the five UPPS-P subscales were associated with substance use, and only UPPS-P “positive urgency” and “negative urgency” were associated with anxiety or depression symptoms. IPTs are also thought to be related to delay discounting, a tendency to devalue future events or rewards (Moreira & Barbosa, 2019), although these constructs sometimes show small or divergent associations (Murphy & Mackillop, 2012; Reynolds et al., 2006).

IPTs also appear to be genetically dissociable. The largest genome-wide association study (**GWAS**) of IPTs to date (Sanchez-Roige et al., 2019) showed only moderate genetic correlations between UPPS-P and BIS subscales. In addition, the UPPS-P “sensation seeking” scale was only weakly genetically correlated with other IPTs, but was instead more strongly genetically correlated with extraversion. Furthermore, UPPS-P “lack of perseverance” was weakly genetically correlated with other IPTs. These findings are consistent with results from twin studies (Gustavson et al., 2019).

To study the multifaceted nature of impulsivity, we used a recently introduced method, genomic structural equation modeling (genomic SEM; Grotzinger et al., 2019), which applies SEM methods to genetic correlations based on GWAS results, using the same techniques as SEM models based on phenotypic correlations. Because all GWAS use the same ancestral genome as a reference, GWAS summary statistics across different traits and participants can be linked through this common reference. Thus, genetic covariances/correlations can be estimated between any pair of GWAS traits, providing both samples were drawn from the same ancestral background (Bulik-Sullivan et al., 2015). Whereas models based on phenotypic or twin correlations require all traits to be measured in the same sample, genomic SEM models do not, greatly expanding the range of traits and models that can be examined. Moreover, because GWAS analyses do not rely on siblings, they are not subject to the same potential biases that arise from assumptions of twin studies (e.g., the equal environments assumption).

The aims of this study were three-fold (**Figure 1**). First, we leveraged data from our GWAS of UPPS-P and BIS (Sanchez-Roige et al., 2019) to examine the latent genetic structure of impulsivity; namely we tested multiple competing hypotheses about whether single or multiple genetic factors are needed to capture the genetic structure of IPTs. The UPPS-P “negative urgency” and “positive urgency” subscales are often considered two facets of a higher-order factor reflecting emotion-based rash action, given their phenotypic similarities (Cyders & Smith, 2007; Littlefield et al., 2016). Similarly, there is some evidence that UPPS-P “lack of premeditation” and “lack of perseverance” might load onto a common factor representing deficits in conscientiousness (Cyders & Smith, 2007; Whiteside & Lynam, 2001). Therefore, we compared models with 5 unique IPT factors to models with 3 or 4 IPT factors by collapsing positive and negative urgency and/or lack of premeditation and perseverance. We also considered whether all facets would load on a single factor. In these analyses, we used published GWAS results from an independent study of extraversion (van den Berg et al., 2016) as an additional indicator of sensation seeking, for several reasons. First, UPPS-P “sensation seeking” was genetically correlated with extraversion in our earlier work (Sanchez-Roige et al., 2019). In addition, sensation seeking has long been conceptualized as a component of extraversion in theories of personality (Costa & McCrae, 1992; Eysenck, 1993) and both constructs are measured using questionnaire that contain some nearly identical items.

**Figure 1:**
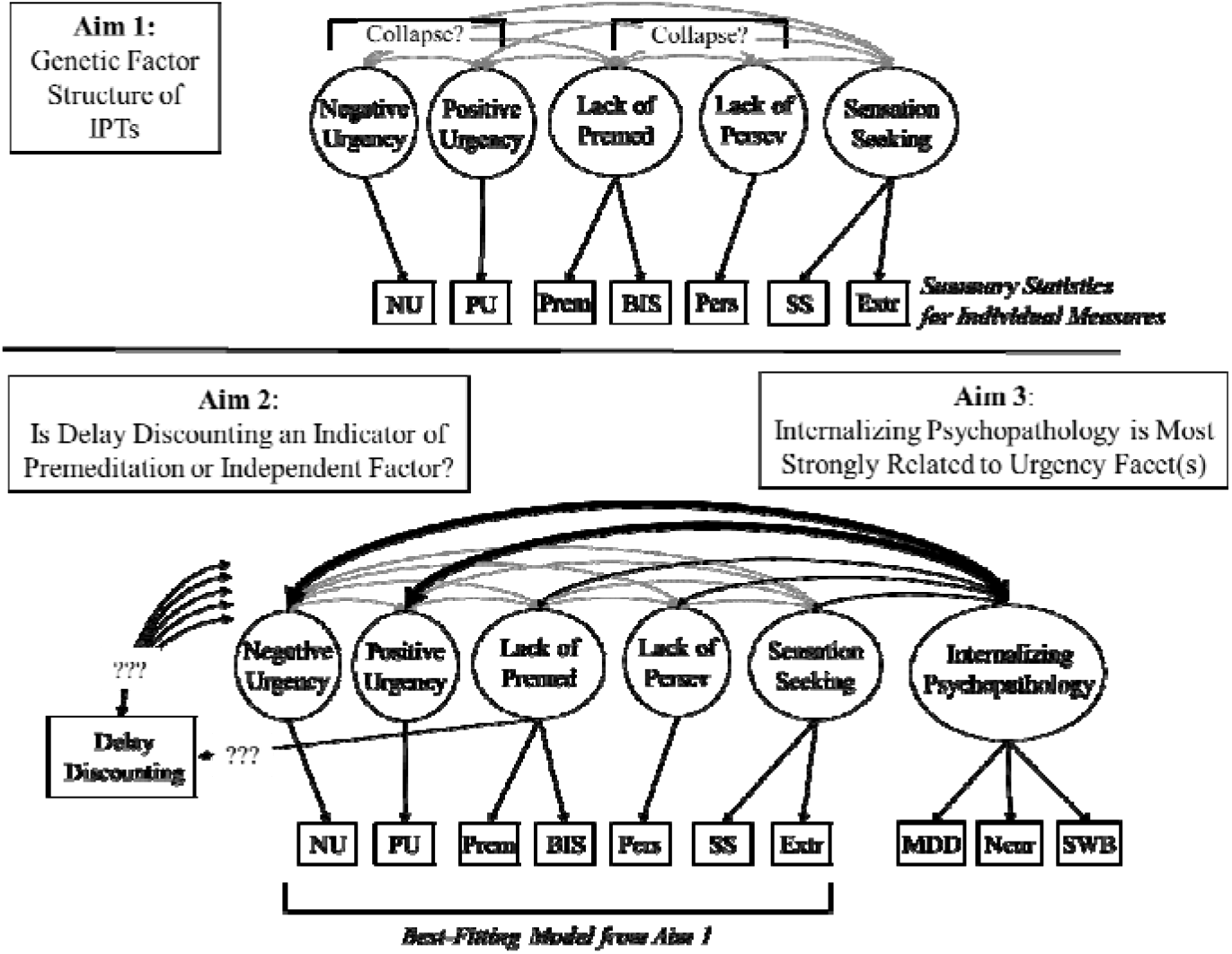
Visual representation of the study aims. In all models, summary statistics for individual impulsive personality traits (IPTs) are represented as squares and latent factors are represented by ovals. Aim 1 was to evaluate whether the 5 IPTs were captured by separate genetic factors or whether certain facets could be collapsed together (e.g., positive and negative urgency and/or lack of perseverance and lack of premeditation, as in Cyders & Smith, 2007). In these models, UPSS-P subscales were initially modeled as separate factors, with the BIS total score loading on the Lack of Premeditation factor. Extraversion was included as a second indicator of sensation seeking to aid in model fit, but this did not affect the pattern of results (see **Supplement**). Aims 2 and 3 were evaluated simultaneously by adding Delay Discounting and Internalizing Psychopathology in the same model to provide the maximum information in the genetic correlation matrix. Aim 2 evaluated whether genetic influences on delay discounting were best modelled as an independent genetic factor, or a facet of lack of premeditation. Aim 3 was to evaluate the hypothesis that a latent factor capturing genetic influences on internalizing psychopathology would be most strongly genetically correlated with IPT genetic factors related to control over emotion-based rash action (positive urgency, negative urgency, or their combination, depending on Aim 1). Genetic correlations among IPTs are shown in grey for simplicity. Lack of Premed = Lack of Premeditation factor; Lack of Persev = Lack of Perseverance factor; NU = UPPS-P negative urgency subscale; PU = UPPS-P positive urgency subscale; Prem = UPPS-P lack of premeditation subscale; BIS = Barratt Impulsiveness Scale; Pers = UPPS-P lack of perseverance subscale; SS = UPPS-P sensation seeking subscale; Extr = Extraversion; MDD = Major depressive disorder; Neur = Neuroticism; SWB = Subjective Wellbeing.

Second, we examined whether delay discounting (**DD**) could be modeled as a common or separate genetic factor. We hypothesized that DD could be an indicator of lack of premeditation, based on previous work showing strong genetic correlations between the two facets (Sanchez-Roige et al., 2019). DD captures the valuation of future versus present rewards whereas lack of premeditation captures the tendency to act without thinking about future consequences. Because both DD and lack of premeditation involve the consideration of future outcomes, they may reflect a common factor. We evaluated this possibility against a model where DD represents a distinct genetic factor that is simply more correlated with lack of premeditation than other IPTs.

Finally, we leveraged published GWAS of phenotypes related to internalizing psychopathology to further inform the genetic structure of IPTs. Previous work has shown that IPTs were associated with internalizing psychopathology, including self-report and diagnostic assessment of major depressive disorder and generalized anxiety disorder (Berg et al., 2015), and that these correlations were driven by shared genetic influences (Gustavson et al., 2019). Thus, we evaluated whether genetic separability among IPTs was accompanied by differential relations to an internalizing psychopathology factor based on data from well-powered published GWAS of major depressive disorder (Howard et al., 2018), neuroticism (Luciano et al., 2018), and subjective-wellbeing (Okbay et al., 2016). Building on previous phenotypic analyses (Berg et al., 2015; Carver & Johnson, 2018), we hypothesized that specific IPTs, particularly those pertaining to emotion regulation (i.e., negative/positive urgency), would be more strongly genetically correlated with internalizing psychopathology than other IPTs, supporting their distinction from other IPTs.

## Method

### Genome-wide association studies

#### IPTs

We used GWAS summary statistics for IPTs from our previously published work (Sanchez-Roige et al., 2019); these association results included measures from the UPPS-P Impulsive Behavior Scale (Cyders, Littlefield, Coffey, & Karyadi, 2014; Whiteside & Lynam, 2001) and the BIS (Patton et al., 1995). The 20-item brief version UPPS-P Impulsive Behavior Scale includes 4 items for each subscale (“lack of premeditation”, “lack of perseverance”, “positive urgency”, “negative urgency”, and “sensation seeking”). Although the 30-item BIS is comprised of three subscales (“attentional”, “motor”, and “nonplanning”), genetic correlations among the subscales were essentially 1.0 (Sanchez-Roige et al., 2019), suggesting that the BIS subscales largely capture a single set of genetic influences related to lack of premeditation. Therefore, we limited our analyses to BIS total score. All research participants included in the analyses were of European ancestry and were research participants from 23andMe, Inc. The final number of research participants included in the analyses range from 21,495 to 22,861. These datasets have been extensively described elsewhere (Sanchez-Roige et al., 2019).

#### Extraversion

Publicly available GWAS summary statistics for extraversion were obtained from a recent meta-analysis of 63,030 individuals of European ancestry (van den Berg et al., 2016). Extraversion was assessed with a harmonized measure across 29 cohorts who administered common measures of extraversion (including the NEO Personality Inventory, NEO Five Factor Inventory, the Eysenck Personality Questionnaire [EPQ], the Eysenck Personality Inventory [EPI], the Reward Dependence scale of Cloninger’s Tridimensional Personality Questionnaire, and the Positive Emotionality scale of the Multidimensional personality Questionnaire). Multiple items from these questionnaires refer to the tendency to enjoy and seek out exciting situations (e.g., “Do you often long for excitement?” [EPI], “Would you do almost anything for a dare?” [EPI], “Do you like plenty of action and excitement around you?” [EPQ]). Although we included extraversion in our initial model of IPTs to aid in model identification, similar models were evaluated without extraversion and its inclusion did not alter the pattern of results (see **Supplement**).

#### Delay discounting

We used GWAS summary statistics for DD from our previous study (Sanchez-Roige et al., 2018), which was based on 23,127 European research participants from 23andMe. Participants completed the 27-item Monetary Choice Questionnaire, a widely used measure of DD (Kirby, Petry, & Bickel, 1999).

#### Internalizing psychopathology

We used publicly available summary statistics for internalizing psychopathology from three independent GWAS: depression (170,756 cases and 329,443 controls; Howard et al., 2018), neuroticism (N=390,278; Luciano et al., 2018), and subjective well-being (N=298,420; Okbay et al., 2016). All individuals were of European ancestry. Depression was assessed in the UK Biobank based on whether an individual had a diagnosis of a depressed mood disorder from linked hospital records or if they answered yes to either of the following questions at any assessment: “Have you ever seen a general practitioner (GP) for nerves, anxiety, tension or depression?” or “Have you ever seen a psychiatrist for nerves, anxiety, tension or depression?”. Neuroticism was also assessed in the UK Biobank with a 12-item version of the Eysenck Personality Inventory-Revised Short Form (Luciano et al., 2018). Subjective well-being was assessed with multiple study-specific measures, although the majority used validated life satisfaction scales such as the Satisfaction with Life Scale or the Geriatric Depression Scale (Okbay et al., 2016). Subjective well-being was reverse scored in these analyses such that higher scores indicate lower well-being (i.e., more internalizing problems).

### Data Analyses

Analyses were conducted using the genomic SEM R package (Grotzinger et al., 2019), a novel statistical method that applies SEM methods to GWAS results. Genomic SEM is an extension of linkage disequilibrium score regression (Bulik-Sullivan et al., 2015), which calculates genetic correlations with any two traits with summary statistics available, provided the samples were drawn from the same ancestral background. Using linkage disequilibrium score regression, genomic SEM computes a full genetic correlation matrix across the set of traits for which GWAS summary statistics are provided, and then estimates the model using this correlation matrix using the lavaan package in R. **Table S1 and Figure 2** display the final genetic correlation matrix for analyses of Aims 2 and 3 that includes all study variables.

**Figure 2:**
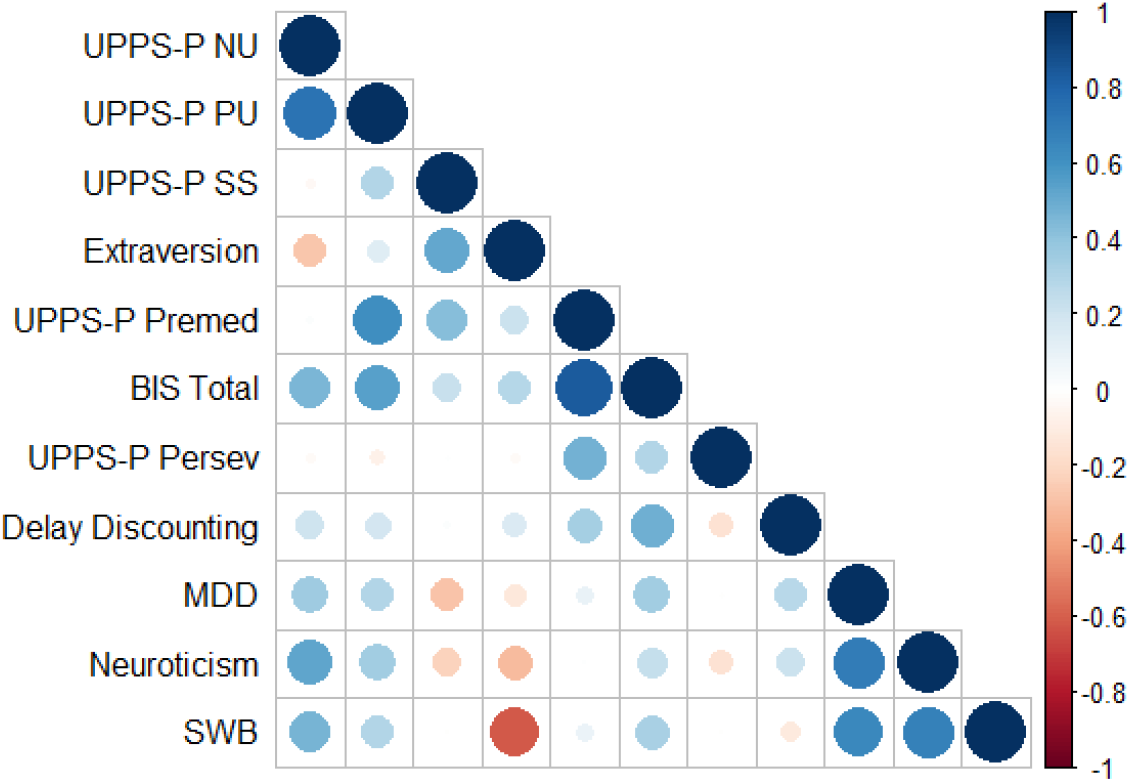
Genetic correlation matrix generated by genomic SEM for Aim 2 and 3 analyses involving all study measures. Matrices for Aim1 (IPTs only) were similar but not identical because each matrix is generated separately in genomic SEM. See supplementary Table S1 for exact *r* values. UPPS-P = UPPS-P Impulsive behavior scale; NU = negative urgency subscale; PU = positive urgency subscale; SS = sensation seeking subscale; Premed = lack of premeditation subscale; BIS = Barratt Impulsiveness Scale; Persev = lack of perseverance subscale; MDD = Major depressive disorder; SWB = Subjective wellbeing (reverse scored). See the online version of the article for the color version of this figure.

Most of the summary statistics included in the analyses were based on overlapping samples (e.g., 23andMe, UK Biobank); however, this method adjusts for sample overlap by estimating a sampling covariance matrix which indexes the extent to which sampling errors of the estimates are associated (Grotzinger et al., 2019).

The R datafiles containing these genomic SEM matrices for all analyses are displayed at the following link (https://osf.io/5x3ft/?view_only=95ee8b4033f54e7e85cfdd2dfe00ae97), alongside R analysis scripts. Supplement **Table S1** also displays the genomic SEM matrix for Aim 3 (which includes all measures examined here).

We applied genomic SEM to test confirmatory factor models that were informed by psychology and psychometric theories (Carver & Johnson, 2018; Cyders & Smith, 2007). We used the default Diagonally Weighted Least Squares (DWLS) estimation method. We used a series of metrics to evaluate the best-fitting confirmatory factor model. Specifically, model fit was determined based on chi-square tests (χ^2^), the Comparative Fit Index (**CFI**) and Akaike Information Criterion (**AIC**). Good-fitting models are expected to have CFI >.95 and smaller AIC values than competing nested models (Hu & Bentler, 1998). Good-fitting models also traditionally have nonsignificant χ^2^ statistics, but because GWAS sample sizes are extremely large, and this statistic is sensitive to sample size, we focused on other fit indices. Significance of individual parameter estimates were established with 95% confidence intervals (95% CIs) and with χ^2^ difference tests (χ^2^_diff_). When fitting models with only 2 indicators, standardized factor loadings were equated to help identify the model. In some cases, “dummy” latent factors were created when we had only a single indicator (e.g., lack of perseverance), with fixed factor loading at 1.0 on their single indicators (and no residual variance). We refer to these as “factors” in the Results, but they should be interpreted as the single indicators that they represent.

Our recent GWAS of IPTs (Sanchez-Roige et al., 2019) indicated that we had power to observe genetic correlations between IPTs and the other traits examined here, and power to detect correlations is typically larger for latent constructs.

## Results

### Genomic Structure Equation Modeling of Impulsivity Facets

We first fitted a genomic SEM model using GWAS data from UPPS-P and BIS subscales, and extraversion. **Table 1** displays model comparisons and **Figure 3** shows the bestfitting model (Model 1; see supplement **Table S2** for 95% confidence intervals), *χ*^2^(9) = 12.52, *p* = .185, CFI = .959, AIC = 50.52. This model included all five factors: Negative Urgency (UPPS-P “negative urgency”), Positive Urgency (UPPS-P “positive urgency”), Lack of Premeditation (UPPS-P “lack of premeditation” and BIS total score), Sensation Seeking (UPPS-P “sensation seeking” and extraversion), and Lack of Perseverance (UPPS-P “lack of perseverance”).

**Table 1:**
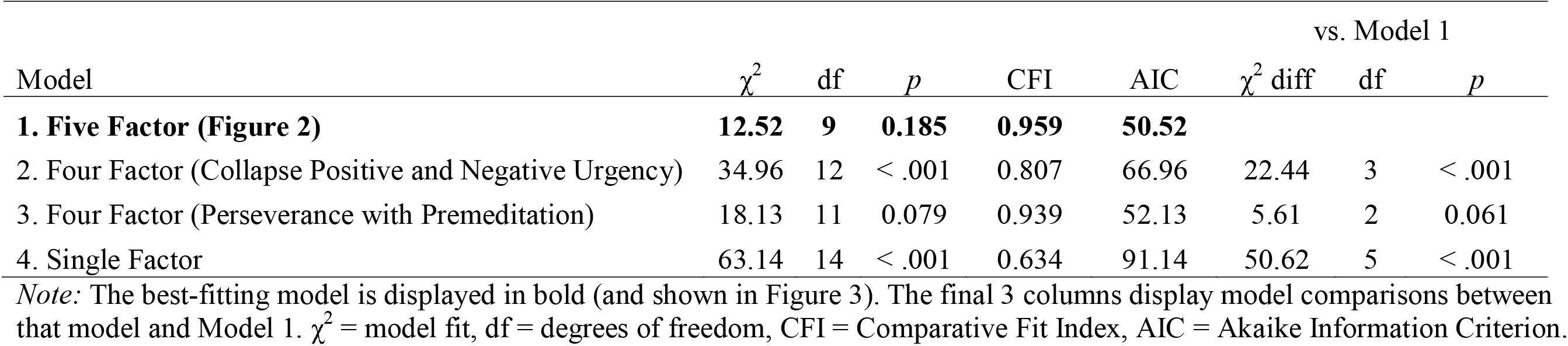
Comparison of Genetic Structure Equation Models of Impulsivity Facets

| Model | $\chi^2$ | df | $p$ | CFI | AIC | vs. Model 1 | | |
| --- | --- | --- | --- | --- | --- | --- | --- | --- |
| | | | | | | $\chi^2$ diff | df | $p$ |
| **1. Five Factor (Figure 2)** | **12.52** | **9** | **0.185** | **0.959** | **50.52** |  |  |  |
| 2. Four Factor (Collapse Positive and Negative Urgency) | 34.96 | 12 | < .001 | 0.807 | 66.96 | 22.44 | 3 | < .001 |
| 3. Four Factor (Perseverance with Premeditation) | 18.13 | 11 | 0.079 | 0.939 | 52.13 | 5.61 | 2 | 0.061 |
| 4. Single Factor | 63.14 | 14 | < .001 | 0.634 | 91.14 | 50.62 | 5 | < .001 |

*Note:* The best-fitting model is displayed in bold (and shown in Figure 3). The final 3 columns display model comparisons between that model and Model 1. $\chi^2$ = model fit, df = degrees of freedom, CFI = Comparative Fit Index, AIC = Akaike Information Criterion.

**Figure 3:**
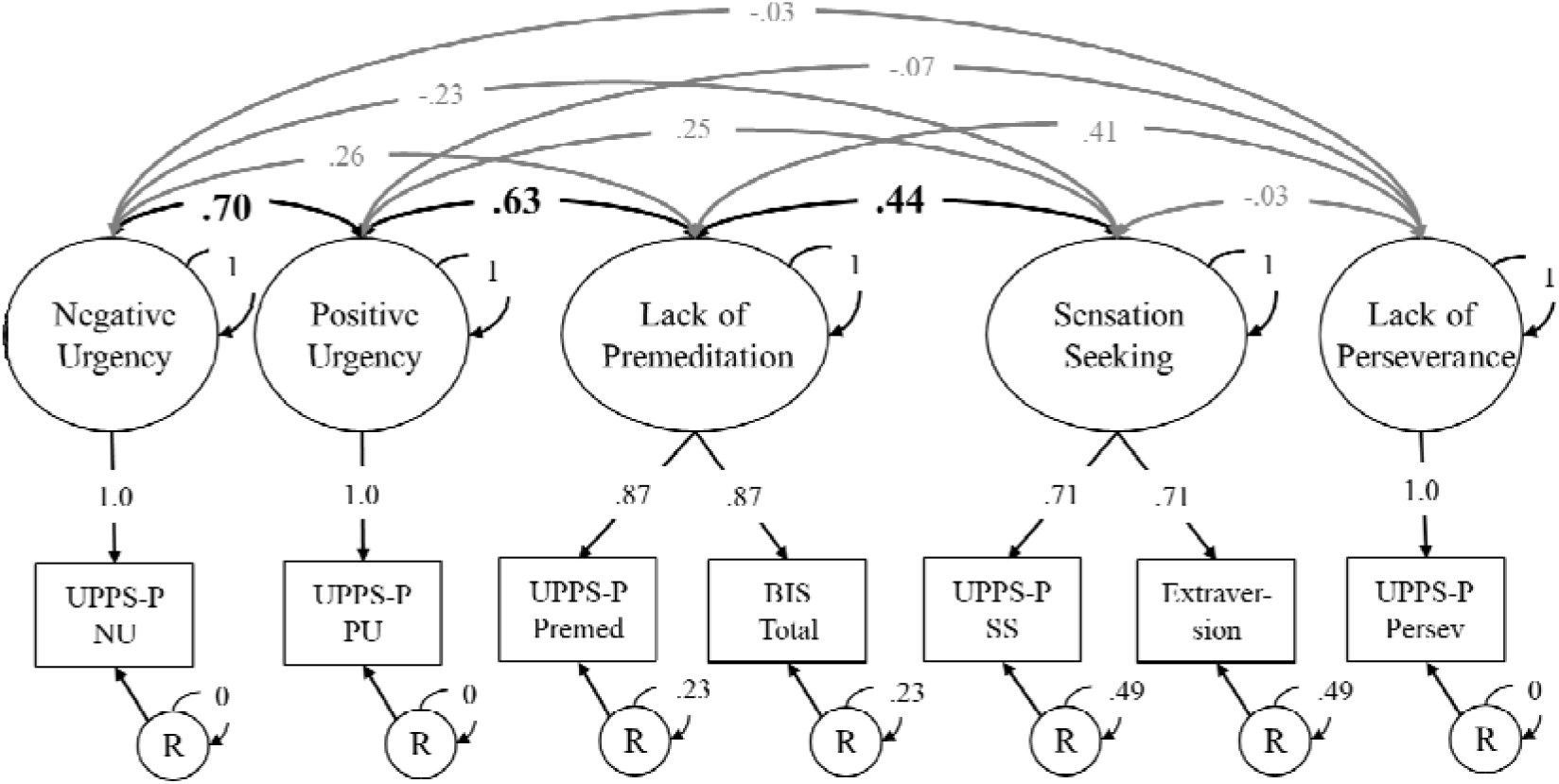
Best-fitting model of the genetic factor structure of impulsivity facets. All individual measures (rectangles) are based on summary statistics from genome-wide association studies. Factor loadings on factors with only two indicators were equated to identify the factor. Factors with only one indicator had factor loadings fixed to 1.0 and residual variances (R) for that indicator fixed to 0. Significant factor loadings and correlations between factors are displayed with bold font and black arrows (based on 95% CIs). Confidence intervals are shown in **Table S2**; confidence intervals were nearly identical to those displayed in **Table 1** and **Table 2** after adding other constructs to the model. All values reflect fully standardized parameter estimates.

We compared Model 1 against models in which UPPS-P “positive urgency” and “negative urgency” subscales were collapsed into a single factor (Model 2), or the BIS total score, UPPS-P “lack of perseverance”, and UPPS-P “lack of premeditation” were collapsed into a single factor (Model 3). Model 2 did not fit the data as well as Model 1, χ^2^_diff_(3) = 22.44, *p* < .001. Although the fit of Model 3 was similar to Model 1, χ^2^_diff_(2) = 5.61, *p* = .061, all other model fit statistics were less favorable (e.g., lower CFI value, higher AIC value, see **Table 1**). Additionally, parallel analyses that did not include extraversion indicated that Model 3 fit worse than Model 1 based on all fit statistics including χ^2^ (see supplement **Table S3 and S4**). Thus, we also rejected Model 3. Finally, Model 4, which was a single-factor model, fit the data very poorly (**Table 1, Model 4**).

### Associations Between Impulsivity and Delay Discounting

We first included a separate DD factor (i.e., with only one indicator), and allowed it to correlate with all other factors in the model. This model, displayed in **Table 2**, showed acceptable fit, *χ^2^*(29) = 154.10, *p* < .001, CFI = .957, AIC = 228.10. As expected, the DD factor was positively genetically correlated with all other IPT factors in the model, although some were not statistically significant (see **Table 2**). Also anticipated, the DD factor was most strongly genetically correlated with the Lack of Premeditation factor, *r* = .47, 95% CI [.03, .91]. However, when we attempted to incorporate DD as a third indicator of the Lack of Premeditation factor, the model fit significantly worse, χ^2^_diff_(5) = 11.71, *p* = .039; overall model fit *χ^2^*(34) = 165.81, *p* < .001, CFI = .955, AIC = 229.81. Thus, DD does not seem to be subsumed as an indicator of lack of premeditation.

**Table 2.** Genetic Correlations Between Impulsivity, Internalizing Psychopathology, and Delay Discounting with 95% Confidence Intervals

| Latent Factor | Indicator | Factor Loading | Genetic Correlations with Other Factors |  |  |  |  |  |
| --- | --- | --- | --- | --- | --- | --- | --- | --- |
|  |  |  | 1 | 2 | 3 | 4 | 5 | 6 |
| 1. Negative Urgency | UPPS-P Negative Urgency | 1 | 1 |  |  |  |  |  |
| 2. Positive Urgency | UPPS-P Positive Urgency | 1 | 0.70<br>[.20, 1.0] | 1 |  |  |  |  |
| 3. Lack of Premeditation | UPPS-P Lack of Premeditation<br>BIS Total Score | 0.87<br>0.87 | 0.26<br>[-.15, .67] | 0.63<br>[.18, 1.0] | 1 |  |  |  |
| 4. Sensation Seeking | UPPS-P Sensation Seeking<br>Extraversion | 0.70<br>0.70 | -0.23<br>[-.60, .14] | 0.24<br>[-.11, .59] | 0.44<br>[.02, .85] | 1 |  |  |
| 5. Lack of Perseverance | UPPS-P Lack of Perseverance | 1 | -0.03<br>[-.42, .35] | -0.06<br>[-.45, .33] | 0.41<br>[-.01, .84] | -0.03<br>[-.31, .26] | 1 |  |
| 6. Internalizing Psychopathology | Major Depressive Disorder<br>Neuroticism<br>Subjective Well-Being | 0.79<br>0.87<br>0.78 | 0.55<br>[.43, .67] | 0.38<br>[.25, .51] | 0.25<br>[.08, .42] | -0.43<br>[-.59, -.27] | -0.10<br>[-.22, .03] | 1 |
| 7. Delay Discounting | Delay Discounting | 1 | 0.20<br>[-.19, .58] | 0.19<br>[-.22, .59] | 0.47<br>[.03, .91] | 0.15<br>[-.18, .47] | -0.16<br>[-.57, .25] | 0.23<br>[.10, .35] |

*Note:* Columns 2 and 3 display the factor loadings from individual measures to latent factors. The remaining columns display the genetic correlations (and corresponding 95% confidence intervals) between latent factors. Factor loadings of 1.0 were fixed to identify these ‘dummy’ latent factors. Factor loadings for both measures in the Lack of Premeditation and Sensation Seeking factors were equated to identify the latent factor. See **Supplementary Figure S1** for a visual depiction of this model.

### Associations Between Impulsivity Facets and Internalizing Psychopathology

As shown in **Table 2**, and as expected, the Internalizing Psychopathology factor was most strongly positively genetically correlated with Negative Urgency, *r* = .55, 95% CI = [.43, .67], and Positive Urgency, *r* = .38, 95% CI = [.25, .51]. The genetic correlation of internalizing with Negative Urgency was significantly stronger than that with Positive Urgency, χ^2^_diff_(1) = 10.93, *p* < .001, but the genetic correlation with Positive Urgency was not significantly stronger than that with Lack of Premeditation, *r* = .25, 95% CI = [.08, .24], χ^2^_diff_(1) = 1.25, *p* = .264. The Sensation Seeking factor was negatively genetically correlated with the Internalizing Psychopathology factor, *r* = −.43, 95% CI = [−.27, −.59], whereas the genetic association with Lack of Perseverance was nonsignificant, *r* = −.10, 95% CI = [−.22, .03]. The Internalizing Psychopathology factor was also associated with Delay Discounting, *r* = .23, 95% CI [.10, .35].

Not surprisingly, this full model confirmed the results of our initial hypothesis regarding the five-factor structure of IPTs. Namely, UPPS-P “positive urgency” and “negative urgency” subscales could not be collapsed into a single factor, χ^2^_diff_(5) = 38.14, *p* < .001, nor could UPPS-P “lack of perseverance” be collapsed into the Lack of Premeditation Factor. Although the model fit was similar, χ^2^_diff_(3) = 0.90, *p* = .825, the CFI and SRMR values were lower and the factor loading for UPPS-P “lack of perseverance” was nonsignificant and in the unexpected direction [−.08].

## Discussion

Numerous studies have examined the phenotypic relationship of IPTs; however, ours is the first to use genomic data to address this question. Our results were consistent with the UPPS-P model, suggesting that positive urgency, negative urgency, lack of premeditation, sensation seeking, and lack of perseverance capture distinct genetic IPTs. DD appeared to represent a unique genetic construct that was only modestly genetically correlated with some of the factors in our models. Of note, each of the genetic factors identified here do not represent individual genes, but rather the contributions of many hundreds/thousands of genetic polymorphisms. This study is also the first to model the genetic structure between IPTs and internalizing psychopathology from unrelated individuals, extending twin research (Gustavson et al., 2019). Internalizing psychopathology was most strongly positively genetically associated with negative and positive urgency, whereas internalizing psychopathology showed a negative genetic correlation with sensation seeking and no genetic correlation with lack of premeditation.

Although our results supported a multi-factor solution of IPTs, it is debatable whether all factors truly represent facets of a single construct of impulsivity. For example, genetic influences on sensation seeking and extraversion loaded highly onto a single genetic factor, with mostly weak or negative genetic correlations with other IPTs. Combined with phenotypic evidence showing that sensation seeking is weakly correlated with other IPTs (MacKillop et al., 2016; Sharma, Markon, & Clark, 2014), and our earlier twin work (Gustavson et al., 2019), this pattern suggests that viewing sensation seeking as a component of impulsivity may represent the “jingle” fallacy (Block, 1995), where different constructs are referred to with the same label. Sensation seeking and impulsivity may instead represent independent processes, or dual systems, with sensation seeking capturing bottom-up reward processing and impulsivity capturing the inability to exert cognitive control (i.e., top-down processing; Shulman, Harden, Chein, & Steinberg, 2016; Steinberg et al., 2008). This result is unsurprising given that the original UPPS study leveraged the Five Factor Model of personality to create the UPPS model (Whiteside & Lynam, 2001), and concluded that the sensation seeking factor corresponded to extraversion. Furthermore, the factor capturing UPPS-P “lack of perseverance” was uncorrelated with all other IPTs, suggesting it is not only genetically distinct from the Lack of Premeditation factor, but may have minimal genetic overlap with other IPTs, consistent with weak phenotypic associations between lack of perseverance and other IPTs in some (Whiteside & Lynam, 2003) but not all studies (Cyders & Smith, 2007; MacKillop et al., 2016). In summary, although our genetic analysis support the current UPPS-P framework that the five IPTs represent five separable constructs (Lynam et al., 2006), it may be more useful to restrict the term “impulsivity” to facets that share some phenotypic and genetic similarities.

On the contrary, the three BIS subscales were so completely genetically correlated that we analyzed them as a single score (as an indicator of the Lack of Premeditation factor). Thus, one framework may be overly inclusive (UPPS-P) and the other may ignore some important aspects of impulsivity (i.e., the BIS does not assess positive or negative urgency).

This work is relevant to our understanding of the role of DD in impulsivity. The theoretical similarity between DD and IPTs has been highlighted across many studies (Moreira & Barbosa, 2019). However, others have argued that low correlations between IPTs and DD measures suggest that these two constructs tap distinct biological mechanisms (Murphy & Mackillip, 2012; Reynolds et al., 2006). Our results that DD was positively, but only weakly-to-moderately, genetically correlated with other IPTs support the latter possibility. DD was most strongly associated with lack of premeditation, consistent with the idea that both constructs relate to the valuation and consideration of future events. Despite their similarities, DD could not be collapsed onto the same latent genetic factor as the other indicators of lack of premeditation, suggesting that DD may be an incremental source of information to include when studying impulsivity.

Finally, our findings confirmed the hypothesis that positive and negative urgency, which are related to emotional control, are more strongly genetically correlated to internalizing psychopathology compared to other IPTs (Carver & Johnson, 2018; Johnson et al., 2013). Moreover, internalizing psychopathology was more strongly positively genetically correlated with negative urgency than positive urgency, consistent with a previous phenotypic metaanalysis (Berg et al., 2015). The positive genetic correlation between lack of premeditation and internalizing psychopathology we observed here was larger than the phenotypic associations with depression and anxiety symptoms found by Berg et al. (2015), suggesting that lack of premeditation may be associated with internalizing psychopathology primarily through genetic influences. In contrast, the Sensation Seeking was negatively genetically correlated with internalizing psychopathology, providing further support for its genetic distinction.

Our findings should be interpreted in the context of the following limitations. First, we demonstrated that certain IPT factors could not be collapsed together on the same factor without a poorer model fit, but some factors only had one indicator and others had only two indicators with equated factor loadings. The latter can contribute to poor model fit to the extent that one measure is a better index of the true latent factor (e.g., UPPS sensation seeking vs. extraversion). Second, IPT measures were based on self-report, and may have different factor structure than laboratory tasks assessing similar constructs (MacKillop et al., 2016; Mallard et al., 2019) However, we anticipate that IPT and task measures will be genetically distinct given their low phenotypic and genetic correspondence (Duckworth & Kern, 2011; Friedman et al., 2019; Sharma et al., 2014). Third, the GWAS data used here reflect ascertainment strategies that may bias our results. For example, the cohorts were generally older, had higher socioeconomic status than the general population, and may have lower than average levels of impulsivity (Sanchez-Roige et al., 2019). In addition, current findings cannot be used to draw inferences about variation among individuals of non-European ancestry, reflecting the underrepresentation of non-Europeans in the field of human genetics. Finally, although the study was based on GWAS of ~20,000-400,000 individuals, many genetic correlations had wide confidence intervals. As sample size for GWAS continues to rapidly increase, this will allow for more precise estimates of associations (and model testing) in future studies.

## Conclusion

Impulsivity is increasingly recognized as a phenotypically heterogeneous construct (Niv et al., 2012), and our genomic SEM analyses provide novel genetic evidence to support this view. The current data support the idea that IPTs tap overlapping but distinct genetic influences. Although sensation seeking and lack of perseverance are considered an impulsivity-related trait within the UPPS-P framework, our data suggest that they are genetically distinct from the other IPTs, consistent with earlier phenotypic observations. DD also appeared to be a distinct genetic factor. Our findings also support the hypothesis that although internalizing psychopathology is positively associated with all impulsivity facets except sensation seeking, this genetic association is most pronounced for IPTs related to the control over negative emotions (Carver & Johnson, 2018). This work demonstrates that large-scale GWAS results can be used to evaluate theoretical models of impulsivity and psychology more broadly.

## Funding

DG was supported by NIH grants DC016977 and HD098859. NPF was supported by NIH grants MH063207, DA046064, DA046413, and DA042742. SSR was supported by the NARSAD Young Investigator Award from the Brain and Behavior Foundation (#27676). SSR and AAP were supported by funds from the California Tobacco-Related Disease Research Program (TRDRP; Grant Number 28IR-0070 and T29KT0526). We would like to thank the research participants and employees of 23andMe for making this work possible.

## Author Contributions

DG and SSR developed the study concept and study design. DG and SSR created correlation matrices from available GWAS summary statistics and conducted all analyses. All authors provided critical revisions and approved the final version of the manuscript for submission.

