## Supplemental Information for "The Latent Genetic Structure of Impulsivity and its Relation to Internalizing Psychopathology"

*Supplement Table S1*

*Genetic Correlations Among all Measures in the Study*

| Measure | 1 | 2 | 3 | 4 | 5 | 6 | 7 | 8 | 9 | 10 | 11 |
|---|---|---|---|---|---|---|---|---|---|---|---|
| 1. UPPS-P Negative Urgency | 1 | | | | | | | | | | |
| 2. UPPS-P Positive Urgency | .74 | 1 | | | | | | | | | |
| 3. UPPS-P Lack of Premeditation | .01 | .62 | 1 | | | | | | | | |
| 4. BIS Total Score | .46 | .54 | .83 | 1 | | | | | | | |
| 5. UPPS-P Sensation Seeking | -.03 | .30 | .42 | .23 | 1 | | | | | | |
| 6. Extraversion | -.28 | .13 | .22 | .28 | .52 | 1 | | | | | |
| 7. UPPS-P Lack of Perseverance | -.03 | -.07 | .47 | .30 | .01 | -.03 | 1 | | | | |
| 8. Major Depressive Disorder | .36 | .29 | .10 | .35 | -.29 | -.13 | -.01 | 1 | | | |
| 9. Neuroticism | .53 | .35 | .00 | .24 | -.22 | -.31 | -.16 | .69 | 1 | | |
| 10. Subjective Well-Being | .46 | .29 | .08 | .33 | .00 | -.61 | .00 | .64 | .68 | 1 | |
| 11. Delay Discounting | .20 | .18 | .33 | .49 | .01 | .15 | -.16 | .27 | .21 | -.11 | 1 |

*Note:* This correlation matrix is generated by genomic SEM and used for analyses addressing Aims 2 and 3 involving all study measures (also displayed in Figure 2 of the main text). The matrix for Aim1 (IPTs only) was similar but not identical because each matrix is generated separately in genomic SEM.

*Supplemental Table S2*

*Aim 1 Best-Fitting Model (with 95% Confidence Intervals)*

| Indicator | Factor Loading | Genetic Correlations with Other Factors 1 | 2 | 3 | 4 | 5 |
|---|---|---|---|---|---|---|
| UPPS-P Negative Urgency | 1 | 1 | | | | |
| UPPS-P Positive Urgency | 1 | 0.71 [.20, 1.0] | 1 | | | |
| UPPS-P Lack of Premeditation | 0.87 | 0.26 [-.15, .67] | 0.63 [.18, 1.0] | 1 | | |
| BIS Total Score | 0.87 | | | | | |
| UPPS-P Sensation Seeking | 0.71 | -0.23 [-.60, .14] | 0.25 [-.11, .60] | 0.44 [.02, .85] | 1 | |
| Extraversion | 0.71 | | | | | |
| UPPS-P Lack of Perseverance | 1 | -0.03 [-.42, .35] | -0.07 [-.46, .32] | 0.41 [-.01, .82] | -0.03 [-.31, .26] | 1 |

*Note:* This model corresponds to Figure 2 in the main text, but also includes 95% confidence intervals around estimates of correlations between latent factors.

*Supplemental Table S3*

*Aim 1 Model Comparisons (Excluding Extraversion)*

| | | | | | | vs. Model 1 | | |
|---|---|---|---|---|---|---|---|---|
| Model | $\chi^2$ | df | *p* | CFI | AIC | $\chi^2$ diff | df | *p* |
| 1. Five Factor (Figure 2 with no Extraversion) | 8.71 | 4 | 0.069 | 0.944 | 42.72 | - | - | - |
| 2. Four Factor (Collapse Positive and Negative Urgency) | 22.07 | 7 | 0.002 | 0.834 | 50.07 | 13.36 | 3 | 0.004 |
| 3. Four Factor (Collapse Lack of Perseverance and Premeditation) | 15.05 | 6 | 0.02 | 0.912 | 45.05 | 6.34 | 2 | 0.042 |
| 4. Single Factor | 42.02 | 9 | < .001 | 0.679 | 66.02 | 33.31 | 5 | < .001 |

*Note*: These models correspond to those displayed in Table 1, except that the summary statistics for extraversion were excluded from the model. Model 1 still provided the best fit to the data.

*Supplemental Table S4*

*Aim 1 Best-Fitting Model (Excluding Extraversion)*

| Factor | Indicator | Factor Loading | Genetic Correlations with Other Factors 1 | 2 | 3 | 4 | 5 |
|---|---|---|---|---|---|---|---|
| 1 | UPPS-P Negative Urgency | 1 | 1 | | | | |
| 2 | UPPS-P Positive Urgency | 1 | 0.71 [.21, 1.0] | 1 | | | |
| 3 | UPPS-P Lack of Premeditation<br>BIS Total Score | 0.88<br>0.88 | 0.26 [-.16, .67] | 0.63 [.18, 1.0] | 1 | | |
| 4 | UPPS-P Sensation Seeking | 1 | -0.03 [-.42, .36] | 0.29 [-.09, .68] | 0.34 [-.10, .79] | 1 | |
| 5 | UPPS-P Lack of Perseverance | 1 | -0.03 [-.42, .36] | -0.07 [-.46, .33] | 0.41 [-.01, .84] | 0.01 [-.32, .34] | 1 |

*Note:* This model corresponds to Figure 3 in the main text, except that the summary statistics for extraversion were excluded from the model.

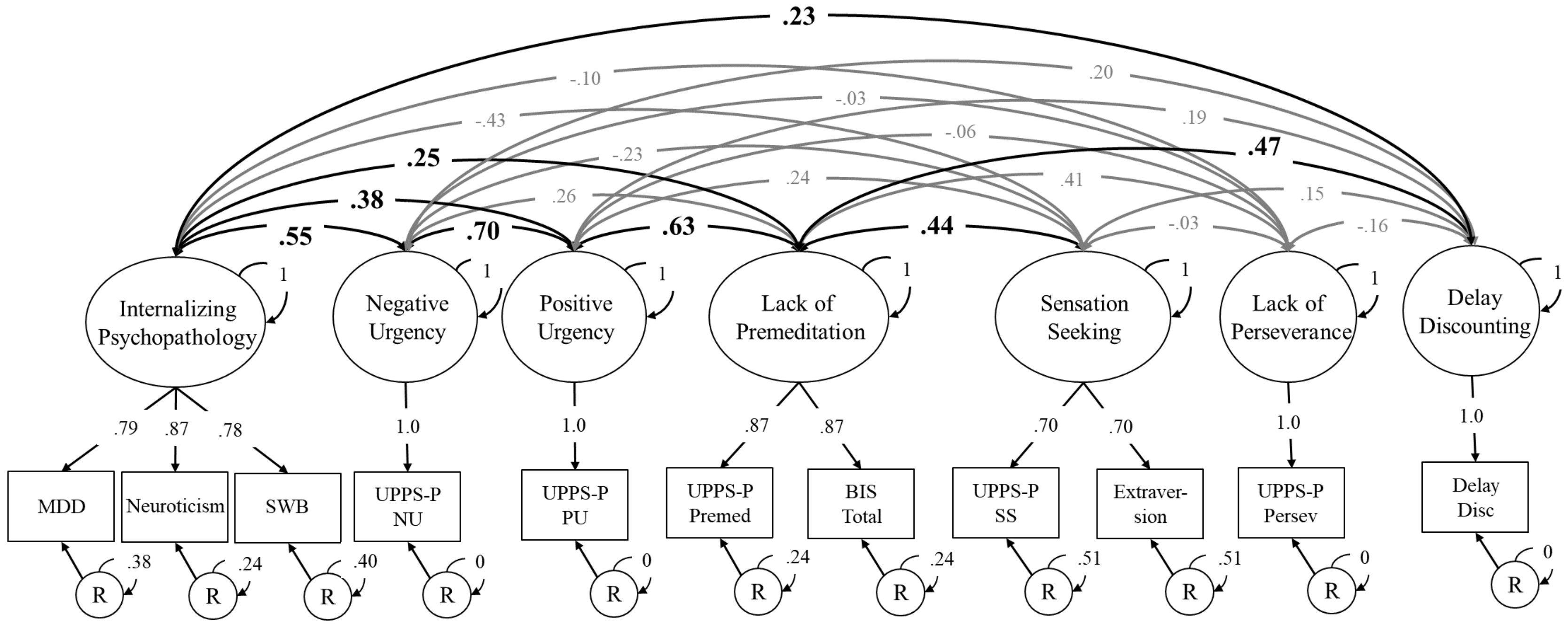


*Figure S1:* This model visually displays the results from **Table 2**, which included all measures. All individual measures (rectangles) are based on summary statistics from genome-wide association studies. Factor loadings on factors with only two indicators were equated to identify the factor. Factors with only one indicator had factor loadings fixed to 1.0 and residual variances (R) for those indicators fixed to 0. Significant factor loadings and correlations between factors are displayed with bold font and black arrows (based on 95% confidence intervals); non-significant correlations are shown with gray lines and regular text. Confidence intervals are shown in **Table 2**. All values reflect fully standardized parameter estimates. NU = negative urgency subscale; PU = positive urgency; Premed = lack of premeditation subscale; BIS = Barratt Impulsiveness Scale; Persev = lack of perseverance subscale; SS = sensation seeking subscale; MDD = Major depressive disorder; SWB = Subjective wellbeing; Delay Disc = Delay Discounting (Monetary Choice Questionnaire).
